# Neural correlates of early sound encoding and their relationship to speech in noise perception

**DOI:** 10.1101/076455

**Authors:** Emily B.J. Coffey, Alexander M.P. Chepesiuk, Sibylle C. Herholz, Sylvain Baillet, Robert J. Zatorre

## Abstract

Speech-in-noise (SIN) perception is a complex cognitive skill that affects social, vocational, and educational activities. Poor SIN ability particularly affects young and elderly populations, yet varies considerably even among healthy young adults with normal hearing. Although SIN skills are known to be influenced by top-down processes that can selectively enhance lower-level sound representations, the complementary role and of feed-forward mechanisms and their relationship to musical training is poorly understood. Using a paradigm that eliminates the main top-down factors that have been implicated in SIN performance, we aimed to better understand how robust encoding of periodicity in the auditory system (as measured by the frequency-following response) contributes to SIN perception. Using magnetoencephalograpy, we found that the strength of encoding at the fundamental frequency in the brainstem, thalamus, and cortex is correlated with SIN accuracy, as was the amplitude of the slower cortical P2 wave, and these enhancements were related to the extent and timing of musicianship. These results are consistent with the hypothesis that basic feed-forward sound encoding affects SIN perception by providing better information to later processing stages, and that modifying this process may be one mechanism through which musical training might enhance the auditory networks that subserve both musical and language functions.

**Highlights:**

– Enhancements in periodic sound encoding are correlated with speech-in-noise ability
– This effect is observed in the absence of contextual cues and task demands
– Better encoding is observed throughout the auditory system and is right-lateralized
– Stronger encoding is related to stronger subsequent secondary auditory cortex activity
– Musicianship is related to both speech-in-noise perception and enhanced MEG signals

## 1.1 Introduction

The ability to decipher speech in the presence of background noise affects people's participation in social, vocational, and educational activities [Anderson and Kraus, 2010]. Understanding the neural bases of good speech-in-noise **(SIN)** perception during development, adulthood, and into old age is both clinically and scientifically important, but it is challenging due to the complexity of the skill, which involves interactions between peripheral hearing and central processing, and because it is affected by life experiences [Anderson et al., 2013a]. Here, we aimed to better understand the neural bases of periodicity coding in the brain to understand its relevance to SIN, and how it might be enhanced by musicianship.

One means of observing the inter-individual differences in how people encode periodic characteristics of sound is the frequency-following response (FFR), an evoked response that is an index of the temporal representation of periodic sound in the brainstem [Chandrasekaran and Kraus, 2010; Skoe and Kraus, 2010], thalamus, and auditory cortex [Coffey et al., 2016b]. Differences in the strength of the fundamental frequency (fO) of the FFR have been linked to SIN perception (e.g. [Kraus and Nicol, 2005]), and FFR-fO strength may also be enhanced by training [Song et al., 2008; Song et al., 2012]. However, enhancements and deficits of neural correlates that are related to SIN perception are most consistently identified either in very challenging listening conditions [Parbery-Clark et al., 2009b] or in the degree of degradation of the signal between quiet and noisy conditions (e.g. [Cunningham et al., 2001; Song et al., 2011]). It is therefore unclear if the observed relationship between the FFR and SIN is due only to better top-down mechanisms such as better stream segregation [Başkent and Gaudrain, 2016] or selective auditory attention [Lehmann and Schonwiesner, 2014; Parbery-Clark et al., 2011; Song et al., 2011]), or if enhanced feed-forward stimulus encoding also plays a role. Although most studies have used electroencephalography (EEG) to record the FFR, we have recently shown that magnetoencephalography (MEG) adds spatial information [Coffey et al., 2016b], which might therefore be useful to investigate the covariation of measures of auditory system enhancement as they pertain to SIN perception.

Musicians are thought to have both enhanced bottom-up [Bidelman and Weiss, 2014; Musacchia et al., 2007] and top-down [Kraus et al., 2012; Strait et al., 2010] processing of sound. Because SIN perception and measures of basic sound encoding are related to musicianship, musical training has been proposed as a means of ameliorating poor SIN performance (reviewed in: [Alain et al., 2014]). Musical training places high demands on sensory, motor, and cognitive processing mechanisms that overlap between music and speech perception, and offers extensive repetition and emotional reward, which could stimulate auditory system enhancements that in turn impact speech processing [Patel, 2014]. Several longitudinal studies support a causal relationship between musical training and SIN skills [Kraus et al., 2014; Slater et al., 2015; Tierney et al., 2013], although it is difficult to maintain full, experimental control over naturalistic training studies [Evans et al., 2014]. A number of cross-sectional studies have also reported a musician advantage in SIN perception [Parbery-Clark et al., 2009a; Parbery-Clark et al., 2009b; Parbery-Clark et al., 2011; Parbery-Clark et al., 2012; Strait et al., 2012; Swaminathan et al., 2015; Zendel and Alain, 2012]; however, other studies have not found significant group differences [Boebinger et al., 2015; Ruggles et al., 2014] or have found the musicianship effect to be dependent upon the specific SIN task variations, such as the degree of information masking [Başkent and Gaudrain, 2016; Swaminathan et al., 2015] or the degree of reliance on pitch cues [Fuller et al., 2014]. It is therefore not clear if there is a consistent musician advantage; if so, if it comes about due to training, self-selection [Schellenberg, 2015] or predispositions [Zatorre, 2013]; and in any case, to which aspects of cognition it is owed: top-down processes such as selective attention and working memory that modulate early levels [Rinne et al., 2008] to filter and temporarily store incoming information [Kraus et al., 2012; Strait and Kraus, 2011], relatively immutable factors such as nonverbal IQ [Boebinger et al., 2015] that might affect multiple cognitive processes, or differences in basic sound encoding (reviewed in [Alain et al., 2014; Anderson and Kraus, 2010; Du et al., 2011], see also [Weiss and Bidelman, 2015]).

In the present study, we aimed to clarify whether robust f0 encoding in the auditory system influences SIN perception in a feed-forward fashion, and whether this variable is related to differences in the neural correlates of later processing stages. If enhanced encoding is partly responsible for better auditory skills because a better quality signal is encoded from incoming sound and passed to higher-order cognitive processes and networks [Irvine, 1986; Musacchia et al., 2008], we would expect that the relationship between SIN and sound encoding would persist even under optimal listening conditions when the system is not challenged, and when the listener's attention is otherwise engaged. In addition to FFR-fO, other cortical potential measures covary with SIN performance, in particular the ERP P2 component (~200 ms post stimulus onset) [Cunningham et al., 2001]), which is also known to be related to speech processing and is sensitive to training effects [Bidelman and Weiss, 2014; Key et al., 2005; Musacchia et al., 2008; Tremblay et al., 2014]. Therefore we would expect enhancements in FFR-f0 to be paralleled by enhancements in the strength of the ERP P2 component, and for each of these measures to be related to SIN accuracy.

To test these hypotheses, we measured SIN perception behaviourally, then simultaneously recorded EEG and MEG data while listeners were presented with a speech sound in quiet as they watched a silent film. We localized the neural origins of the FFR-fO and the P2 and examined their spatial and statistical relationships to each other, and spatial relationships to preceding and following ERP components. We then evaluated the relationships between FFR-f0 components from each main auditory structure and SIN accuracy, and compared each with measures of musical experience. Collectively, these data help us to understand the neural basis of inter-individual differences in SIN perception.

## 2. Methods and materials

The experimental procedures concerning the MEG and (single channel, Cz) EEG recordings of the brain's response to the speech syllable /da/, and much of the pre-processing, have previously been reported in the context of different research questions and will be discussed only briefly here (please see [Coffey et al., 2016b] 'Methods' for details). The correlations between FFR-f0 strength and musicianship that are included in the summary of musical enhancements in Table 1 have been been reported in Coffey et al.; all other findings have not been reported previously.

**Table 1.**
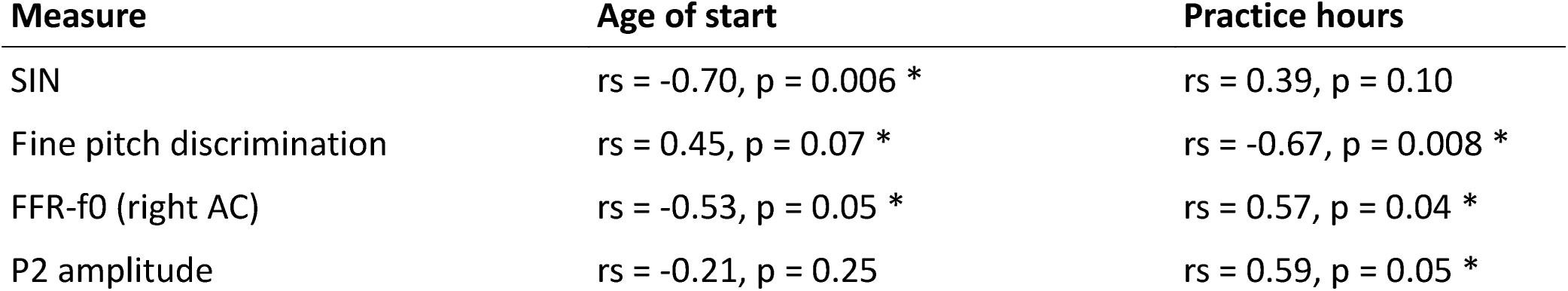
Summary of evidence for musicianship-related behavioural and neurophysiological enhancements (N=12). Asterisks(*) indicate significant rank correlations (alpha < 0.05, one tailed). In general, earlier start ages and a larger number of practice hours are associated with enhancements, suggesting an influence of musical training. Note that lower fine pitch discrimination scores indicate better performance; therefore correlations in opposite directions are expected.

### 2.1. Participants

Data from the same twenty neurologically healthy young adults included in the previous study [Coffey et al., 2016b] were included in this study (mean age: 25.7 years; SD= 4.2; 12 female; all were right-handed and had normal or corrected-to-normal vision;<= 25dB hearing level thresholds for frequencies between 500 Hz and 4,000 Hz assessed by pure-tone audiometry; and no history of neurological disorders). Informed consent was obtained and all experimental procedures were approved by the Montreal Neurological Institute Research Ethics Board.

### 2.2. Speech-in-noise assessment

SIN was measured using a custom computerized implementation of the hearing in noise test (HINT; [Nilsson, 1994]) that allowed us to obtain a relative measure of SIN ability using a portable computer, without specialized equipment. In the standard HINT task, speech-spectrum noise is presented at a fixed level and sentences are varied in a staircase procedure to obtain a SIN perceptual threshold [Nilsson, 1994]. Our modified HINT task used a subset of the same sentence lists [Bench J et al., 1979] and speech-spectrum noise, but presented thirty sentences in three empirically determined difficulty levels in randomized order: easy (2 dB SNR; i.e. target speech was 2 dB louder than noise), medium (-2 dB SNR), and difficult (-6 dB SNR). The sentences and noise were combined using sound processing software (Audacity, version 1.3.14-beta, http://audacity.sourceforge.net/; 44100Hz sampling frequency). Stimuli were presented binaurally via headphones (JVC HA-M5X) with the noise adjusted to a loud but not uncomfortable sound level on pilot subjects (~75 db SPL) and thereafter held constant. SIN accuracy for each difficulty level was calculated as the proportion of words correctly repeated back to the experimenter, and no verbal or visual feedback was given. Values were averaged across difficulty levels to obtain a mean accuracy score.

### 2.3. Fine pitch discrimination

Fine pitch discrimination thresholds were measured as described in [Coffey et al., 2016b], using a two-interval forced-choice task and a two-down one-up rule to estimate the threshold at 79% correct point on the psychometric curve [Levitt, 1971]. The adaptive procedure was stopped after 15 reversals and the geometric mean of the last eight trials was recorded. Thresholds were derived from the average of five task repetitions.

### 2.4 Stimulus presentation

The stimulus for the MEG/EEG recordings was a 120-ms synthesized speech syllable (Ida/) with a fundamental frequency in the sustained vowel portion of 98 Hz. The stimulus was presented binaurally at 80dB SPL, ~14,000 times in alternating polarity, through Etymotic ER-3A insert earphones with foam tips (Etymotic Research). For five subjects, ~11,000 epochs were collected due to time constraints. Stimulus onset synchrony (SOA) was randomly selected between 195 and 205ms from a normal distribution. A separate run was collected of ~600 stimulus repetitions spaced ~5ooms apart, to record later waves of the slower cortical responses. To control for attention and reduce fidgeting, a silent wildlife documentary (Yellowstone: Battle for Life, BBC, 2009) was projected onto a screen at a comfortable distance from the subject’s face.

### 2.5. Neurophysiological recording and preprocessing

Two hundred and seventy-four channels of MEG (axial gradiometers), one channel of EEG data (Cz, 10-20 International System, averaged mastoid references), EOG and ECG, and one audio channel were simultaneously acquired using a CTF MEG System and its in-built EEG system (Omega 275, CTF Systems Inc.). All data were sampled at 12 kHz. Data preprocessing was performed with Brainstorm [Tadel et al., 2011] and using custom Matlab scripts (The Mathworks Inc., MA, USA) as described in [Coffey et al., 2016b].

### 2.6. FFR correlates of SIN accuracy

FFR-f0 strength was extracted from regions of interest (ROls) in the auditory system (AC: auditory cortex, MGB: medial geniculate body of the thalamus, IC: inferior colliculus and CN: cochlear nucleus) using the MEG distributed source modelling approach described previously (see 'Methods: Region of interest spectra from distributed source modelling' in [Coffey et al., 2016b]). We first evaluated correlations between SIN accuracy scores and FFR-f0 strength averaged across bilateral pairs of structures, using Spearman's rho (rs; one-tailed). Non-parametric statistics were used throughout as FFR-fO measures were generally not normally distributed (using Shapiro-Wilk's parametric hypothesis test of composite normality, the null hypothesis was rejected for AC, CN and IC bilateral averages). The EEG equivalent of the FFR-fO was also computed for comparison of sensitivity to behavioural measures. Correlations were computed between SIN accuracy and the left and right auditory cortex ROls separately, as a lateralization effect in FFR-f0 strength and its relationship to measures of musicianship and fine pitch discrimination had been observed previously (see Fig.5c-e in [Coffey et al., 2016b]). We tested for a stronger correlation on the right than left side using Fisher's r-to-Z transformation (one-tailed, alpha= 0.05).

### 2.7 Later cortical evoked responses

Event-related potentials (ERPs) within the 2-40 Hz band-pass filtered single-channel EEG data were obtained in order to establish a connection between previous FFR-ERP research that showed SIN sensitivity at ERP components P2 and N2 [Cunningham et al., 2001] and the MEG data. We did not observe a clear N2 from all subjects. We therefore took the amplitude of only P2 as a measure; this simpler metric also allowed for a more straightforward comparison to and interpretation of the MEG equivalent. A researcher who was blinded to the subjects' FFR-f0 amplitudes and behavioural results at the time of measurement selected P2 wave peaks individually on ERP waves averaged across epochs for each subject (cortically processed; 2-40Hz with −50 to 0 ms DC baseline correction; P2 was considered to be the strongest positive deflection within a ~40 ms window centred on the group grand average P2 at 183 ms). Amplitudes of these custom peaks were then correlated with SIN accuracy and FFR-f0 strength.

MEG evoked response fields (ERFs) on simultaneously recorded data were obtained in order to extend this work using distributed source modelling. The EEG cortical evoked response complex (ERP) elicited by the speech syllable /da/ consists of two positive waves at about 50-90 ms ('Pl') and between 170 and 200 ms ('P2' or 'Pl prime') and two negative waves at about 110 ms ('Nl') and after 200 ms ('N2' or Nl prime') (reviewed in [Key et al., 2005]; see also Cunningham et al., Fig 6 [Cunningham et al., 2001] and Musacchia et al., Fig 2. [Musacchia et al., 2008]). For the purposes of this study we identified wave peaks in the ERP and ERF average at the group level for the SIN-sensitive P2 peak (183 ms), and at the earlier Pl component that has a well-known physiological origin in order to confirm the quality of data and validity of the analysis (60 ms; see Figure 3a,c).

### 2.8. Origins of later cortical ERP components

To confirm that the MEG data could be used to localize areas that showed above-baseline activity at the group level, and to observe the origins of the SIN-sensitive P2 wave in relation to preceding and following ERP waves, we first computed cortical volume MNE models based on each subject's Tl-weighted MRI scan in which the orientation of sources was uncontrained, but their location was constrained within the volume encompassed by the cortical surface. These models were normalized to the baseline period (-50 to Oms). We exported 10 ms time windows around each peak of interest (mean-rectified signal amplitude) and for the baseline (-50 to Oms) for statistical analysis in the neuroimaging software package FSL [Jenkinson et al., 2012; Smith et al., 2004]. These source volume maps were co-registered to the subject's high-resolution Tl anatomical MRI scan (FLIRT, 6 parameter linear transformation), and then to the 2mm MNl152 template (12 parameter linear transformation, [Evans et al., 2012]). Normalized difference images were created by subtracting the baseline images from those of the peaks of interest and calculating z-scores within each image (Pl> Baseline, P2 > Baseline). Permutation testing was used to reveal locations where the magnetic signal was greater during peaks of interest as compared with baseline (non-parametric one-sample t-test [Winkler et al., 2014]; 10,000 permutations). The family-wise error rate was controlled using threshold-free cluster enhancement as implemented in FSL (p < 0.01), after applying a cortical mask of the MNI 152 template with the brainstem and cerebellum removed (these latter structures were not included in the MEG source model).

### 2.9. Comodulation of low and high frequency activity

We considered the spatial relationship between FFR-f0 generators and the source of the SIN-sensitive P2 wave by inspecting the FFR-f0 > Baseline and P2 > Baseline maps in the MEG data, and calculated Spearman's correlations between the FFR-f0 strength from each auditory cortex ROI (MEG) and the amplitude of the P2 wave measured with EEG.

### 2.10 Musicianship enhancements

Within a subset of 12 subjects who reported varying levels of musical experience, we assessed correlations between SIN accuracy and total music practice hours and age of training start, as obtained by self report using the Montreal Music History Questionnaire [Coffey et al., 2011]. We then evaluated the relationship between P2 amplitude in the EEG recording and musicianship, and between fine pitch discrimination skills and musicianship.

## 3. Results

### 3.1. Behavioural scores

The mean accuracy score for the least challenging speech-in-noise condition (2 dB dB difference between speech and noise levels) was 93.5% (SD= 8.0), for the medium difficulty (−2 dB), accuracy was 80.0% (SD= 14.0), and for the most difficult condition (−6 dB), mean accuracy was 33.0 % (SD= 12.0). We averaged only the medium and difficult condition for correlation with neurophyiological measures due to a ceiling effect in the easy condition. The mean averaged SIN score was 56.3% (SD= 12.0). Subjects with finer pitch discrimination ability had statistically better SIN accuracy (one-tailed rs= −0.47, p = 0.018).

### 3.2. MEG FFR-fO strength is related to SIN at each level of the auditory system

As described by Coffey et al., MEG reveals FFR activity from auditory cortex, as well as brainstem and thalamus [Coffey et al., 2016b]. Relationships between FFR values measured via MEG from each ROI (averaged across left and right pairs) and SIN accuracy scores are presented in Figure 1c-f. A positive correlation between SIN accuracy and FFR-f0 strength was found at each of the four structures tested, statistically significant in all but the inferior colliculus, where a similar trend was nonetheless noted. We did not find evidence of a relationship between the EEG-derived FFR-f0 and SIN accuracy (Figure 1g), nor did a relationship appear with the inclusion of age as a covariate (rs= 0.08, p = 0.37).

**Figure 1.**
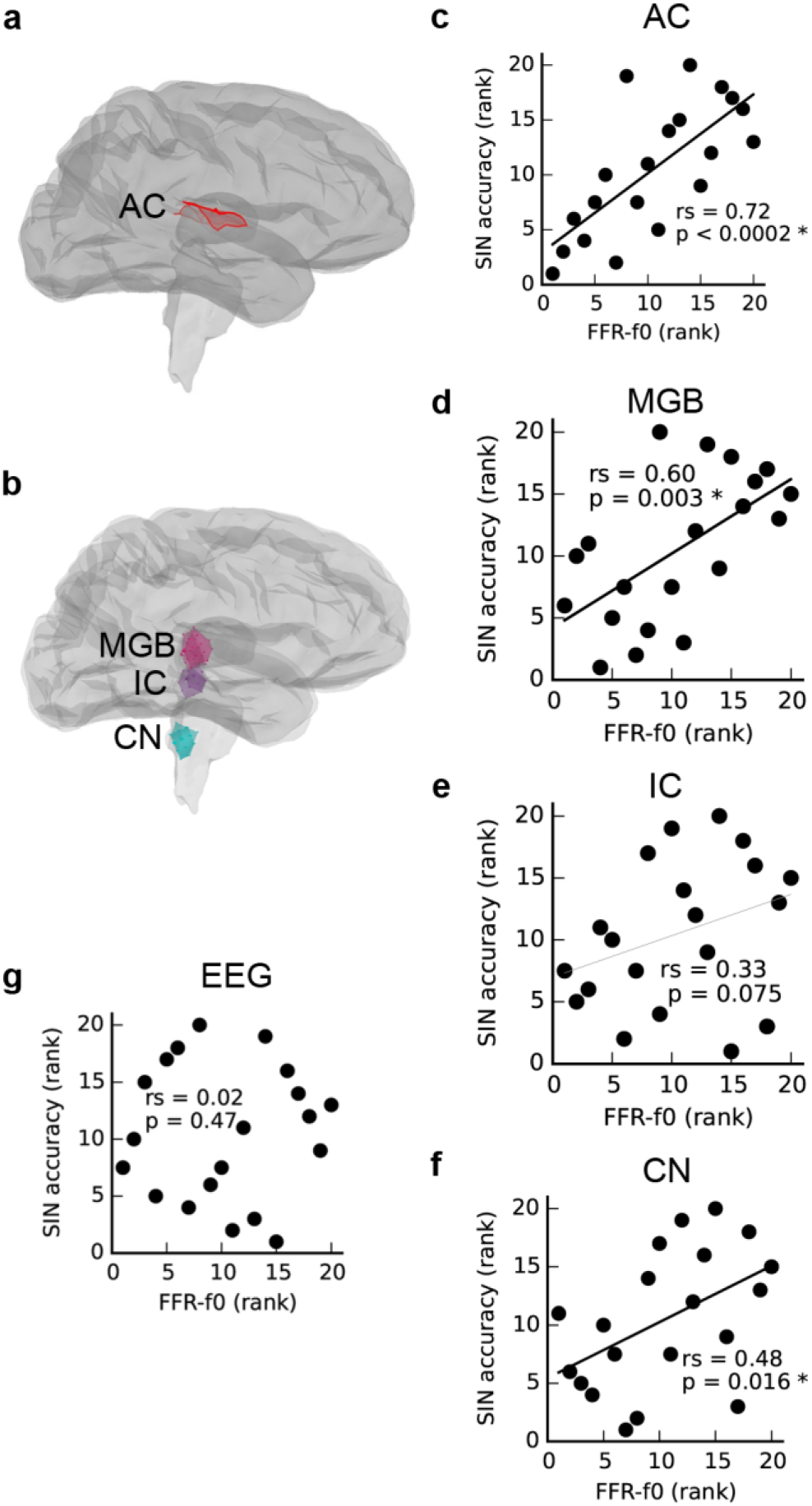
Correlations between FFR-fO strength and speech-in-noise accuracy (SIN) within regions of interest (ROls) in the auditory cortex (a,d), and subcortical areas (b,d-f) as measured with magnetoencephalography (MEG) suggest that better SIN performance is related to better periodicity encoding throughout the auditory system. The FFR measured using electroencephalography at the vertex (Cz) is shown for comparison in (g). AC= auditory cortex, MGB = medial geniculate body, IC= inferior colliculus, CN = cochlear nucleus. Correlations are calculated using Spearman's rho (rs).

### 3.3. The relationship between SIN and cortical FFR-fO is lateralized

The relationship between SIN and FFR-fO strength from auditory cortical ROls in each hemisphere is depicted in Figure 2. SIN accuracy was related to the strength of the FFR-f0 in both hemispheres, but was numerically larger on the right. We directly compared the strength of these correlations using Fisher's r-to-Z–transformation (two-tailed), and found it to be stronger in the right hemisphere (Z = −3.12, p = 0.002; the correlation between the FFR-fO strength across two hemispheres, which is used for statistical comparison of correlation strength, was rs= 0.89).

**Figure 2.**
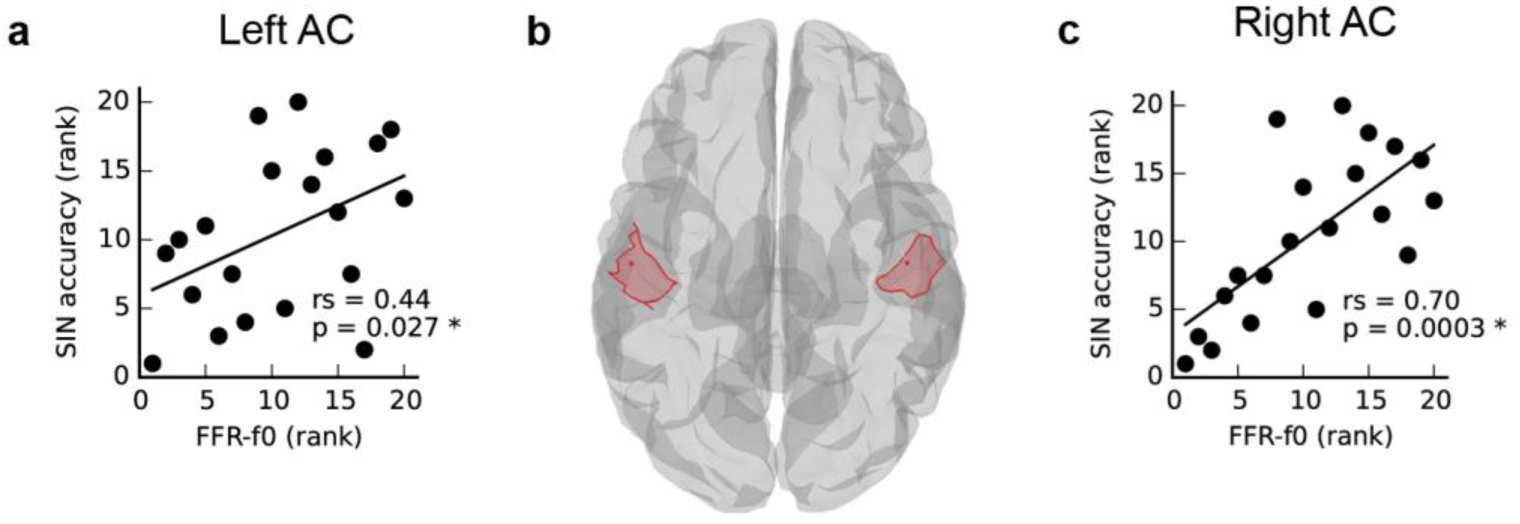
Asymmetry in the relationship between speech-in-noise (SIN) and cortical FFR-fO representation (a,c) within left and right hemisphere auditory cortex ROls, illustrated in (b). Although a positive relationship between FFR-fO strength and SIN is found in each hemisphere, it is significantly stronger in the right hemisphere (Z = −3.118, p = 0.002, two-tailed).

### 3.4. Origins of later cortical ERP components

We confirmed that the MEG data analysis used here is suitable for localizing temporal lobe auditory activity at the group level using Pl, which was known to originate in the primary auditory areas bilaterally (Figure 3d). Note that while we had selected the earliest maximum in the Pl wave in the EEG signal in order to capture primary auditory cortex activity (mean latency: 60 ms), the peak energy in the MEG signal is slightly later (~15 ms); nonetheless, visual inspection of the same analysis performed on a 10 ms window centred on 75 ms indicates that this analysis is not sensitive to minor variations in Pl window selection. The mean latency of P2, the second prominent positive EEG wave, was 183 ms (SD= 11 ms), and its mean amplitude was 4.1 uV (SD= 1.7). We confirmed, as previously reported by Cunningham et al. using a pediatric sample [Cunningham et al., 2001], that P2 amplitude was related to SIN accuracy in the current sample (Figure 3b) thus providing a basis for further investigating FFR-f0 and P2 relationships. As expected, Pl amplitude was not related to SIN accuracy (rs= 0.11, p = 0.33). We then identified the sources of the P2 wave, which proved to be relatively more anterior, and right-lateralized (Figure 3d; coloured areas depict significant clusters corrected for multiple comparisons; maps are thresholded to best expose the areas of strongest signal).

**Figure 3.**
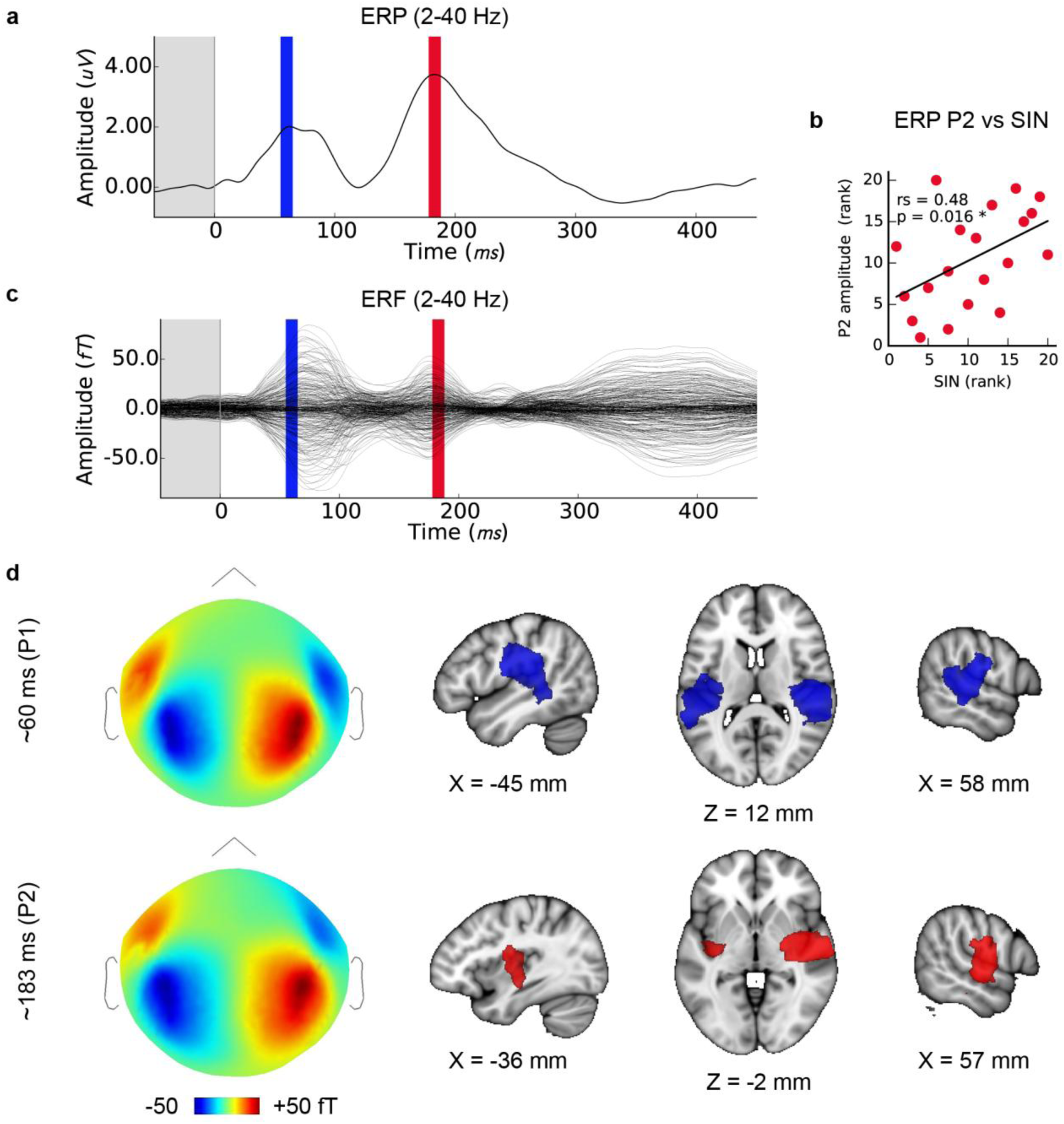
Later cortical evoked responses and their origins. (a) Time courses of the lower frequency evoked response potentials (ERPs) from EEG data with the time windows used for MEG source analysis marked (Pl: blue, P2: red), and (c) evoked response fields (ERFs) from simultaneously recorded MEG data. Each is averaged over subjects (N=20). (b) The amplitude of the P2 ERP wave peak (red) correlates with SIN accuracy. (d) Group-level MEG topographies (left, strength and polarity is indicated in the colour bar below) and source analyses of Pl and P2 component origins using (1mm MNI space; cluster threshold; p < 0.005).

### 3.5. Low and high frequency activity covary

Right but not left AC FFR-f0 strength was significantly related to P2 amplitude (Figure 4c,e). For completeness, we also calculated the correlation between the EEG FFR-f0 and P2 amplitude but it was not significant: rs= 0.23, p = 0.16). The magnetic equivalent of the P2 wave overlapped considerably with the FFR-fO regions using a corrected significance-based threshold of p < 0.05. However, inspection of the centroid of each map showed that whereas the FFR-f0 sources were distributed in the posterior section of the superior temporal gyrus, the P2 wave's foci were more anterior (Figure 4d).

**Figure 4.**
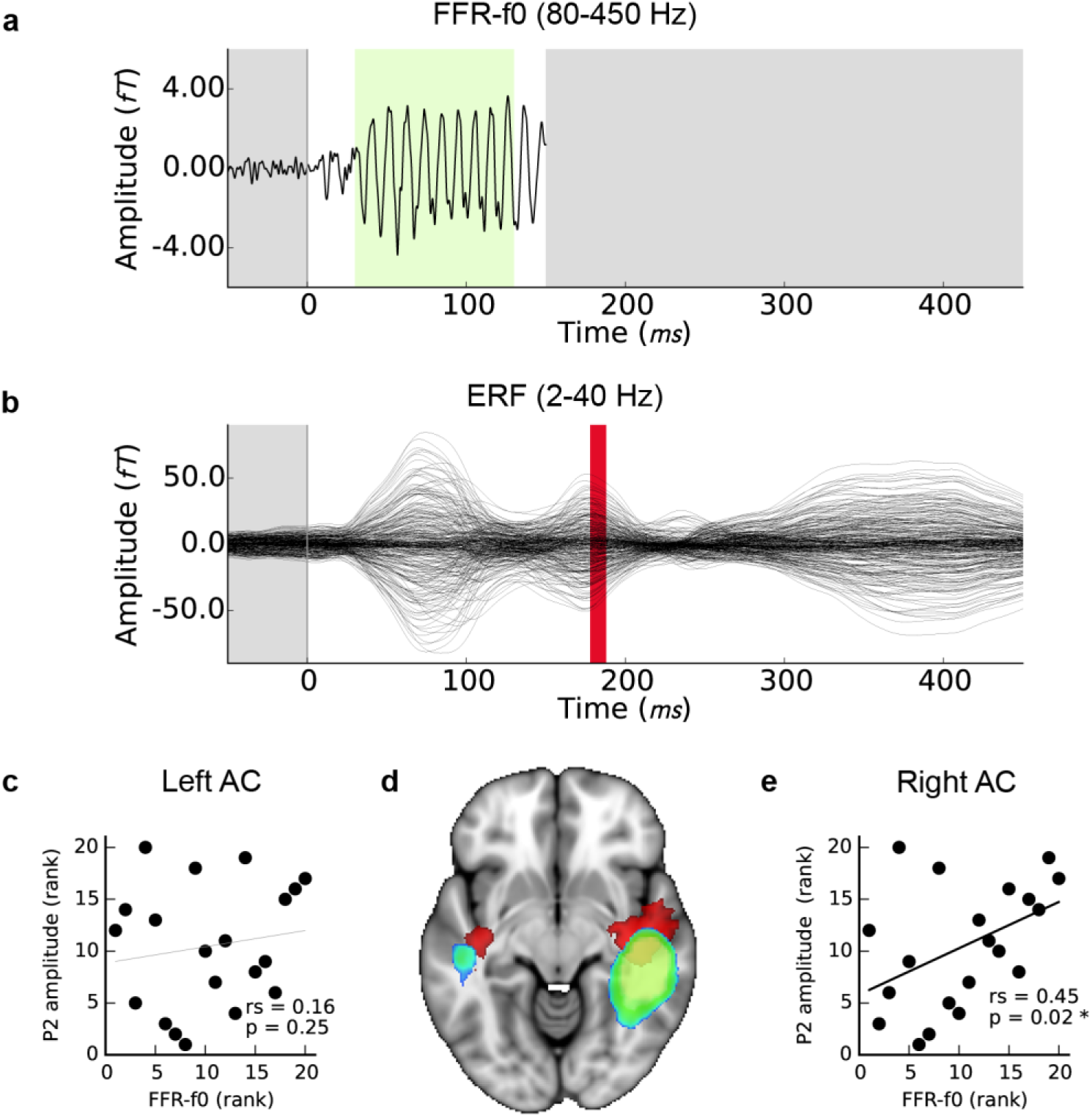
Relationship between FFR-fO strength (blue-green) and P2 amplitude (red). (a) Time course of the FFR-fO response, single channel (as presented in Coffey et al. 2016; 80-450Hz bandpass filtered; −50 to 150 ms window). The green portion indicates period over which FFR-fO strength is calculated. (b) ERF over the same time period; these signals are separated by frequency band (2-40 Hz bandpass filtered) but P2 occurs after the FFR. Correlations between the FFR-fO strength from the cortical ROls as measured using MEG and the ERP strength at P2 as measured using EEG are shown for (c) the left and (e) the right auditory cortex. (d) Illustrates the relationship of the cortical origins of each signal (1mm MNI space; cluster threshold; p < 0.005).

### 3.6. Measures of musicianship

Twelve out of 20 subjects reported some level of musical training. We previously showed that FFR-fO strength in the right AC is related to hours of musical training and age of training start in this sample [Coffey et al., 2016b]. Here, we first confirmed that the correlation between AC FFR-fO strength and SIN scores that we observed within the whole group was present within both the musician subgroup (averaged AC ROls: rs= 0.67, p = 0.012) and the non-musical subgroup (averaged AC ROls: rs= 0.86, p = 0.005). Musical enhancements in behavioural and neurophysiological measures reported in this study are summarized in Table 1. Among those with musical experience, earlier start ages significantly correlated with better SIN scores, but the correlation between total practice hours and SIN showed only a nonsignificant trend. P2 wave amplitude correlated with practice hours, but not the age of start. We also found a significant correlation between hours of musical training and fine pitch discrimination ability, as expected, and a non-significant trend between start age and fine pitch discrimination ability.

## 4. Discussion

In this study, we aimed to clarify whether individual differences observed in fundamental frequency (fO) encoding in the auditory system of normal-hearing adults [Coffey et al., 2016a; Ruggles et al., 2012] influence speech-in-noise (SIN) perception. Towards this end, we measured the neural responses in two frequency bands in order to isolate the higher-frequency FFR (80-450 Hz), and the lower-frequency cortical responses (2-40Hz), and we compared their strength, spatial origins, and relationships to behavioural and musical experience measures.

We first showed that the strength of the MEG-based FFR-fO attributed to each level of the ascending auditory neuraxis, including the auditory cortex in each hemisphere, is related to SIN accuracy (Figure lc-f), suggesting that basic periodic encoding is enhanced throughout the auditory system in people with better ability to perceive speech under challenging noise conditions. Importantly, the only similarity between the conditions under which the neurophysiological measurement was made (i.e. passive listening in silence) and the behavioural measure (i.e. deciphering speech in sentences, similar to the clinical HINT task [Nilsson, 1994]) was the presentation of an auditory speech stimulus. Naturalistic speech-in-noise situations offer multiple cues, many of them redundant, that can result in flexibility in how the task is solved. This is true to a lesser degree of clinical tests of SIN (including the HINT variant used here), which approximate the naturalistic experience of perceiving speech masked by sound, but lack factors that are important in real-life conversations such as familiarity with the talker [Nygaard et al., 1994; Souza et al., 2013], visual cues [Zion Golumbic et al., 2013], and context- and listener-dependent adaptations of the speaker [Lombard, 1911]. Tasks that have been used to study the neural correlates of SIN perception range in cue-richness from natural language comprehension in daily life, to intermediate tasks like the sentence-in-noise and word-in-noise measures, to discrimination of phonemes from among a restricted set of possibilities [Du et al., 2014], and finally to passively listening to single sounds with or without masking noise. Because the effects of variations in paradigm design are unclear [Wilson et al., 2007] even in the clinical measures (for example the HINT may be presented with or without spatial cues [Nilsson, 1994]), this diversity may be contributing to confusion in SIN literature, for example regarding the possibility of a using musical training to improve SIN skills. A systematic analysis of SIN task requirements and differences in their neural correlates may be helpful to clarify these issues (e.g. by using task decomposition [Coffey and Herholz, 2013]). According to such an approach, our neurophysiological recording paradigm did not explicitly engage auditory working memory, it did not offer more than one stream of information, and it did not require attention, which instead was directed to a silent film. Because we have eliminated cues that might be used by these higher-level processes that are known to affect SIN, either by enhancing the incoming signal (e.g. [Lehmann and Schonwiesner, 2014]) or at later linguistic/cognitive processing stages, any residual relationship between basic sound encoding and scores on a high-level SIN task is most parsimoniously explained by the benefits of better lower-level sound encoding. Thus we believe that the correlations we observed at brainstem, thalamic and cortical levels are best interpreted as reflecting these low-level enhancements.

Amplitude differences in the P2 cortical component and in FFR-f0 strength have been related to SIN perception differences between normal and learning-disordered children when recorded in noise [Cunningham et al., 2001]. P2 amplitude also appears to be stronger in musicians [Bidelman and Weiss, 2014; Kuriki et al., 2006; Shahin et al., 2003] (though this has not been observed in all studies [Musacchia et al., 2008]). P2, along with the Pl and the intervening negativity, is affected by stimulus parameters including frequency, location, duration, intensity, and presence of noise (reviewed in [Alain et al., 2013]; see also [Ross and Fujioka, 2016]), is affected by short-term training [Lappe et al., 2011; Tremblay et al., 2014], and is correlated with language–related performance measures such as categorical speech perception [Bidelman and Weiss, 2014]. The findings suggest that the underlying processes are critical to sound representation generally, though the nature and roles of component processes represented in the P2 and their relationships to oscillatory brain networks are still being clarified (e.g. [Ross et al., 2012; Ross and Fujioka, 2016]).

We found that variability in P2 amplitude correlated with inter-individual differences in SIN ability (Figure 3b) and in FFR-fO strength (Figure 4c,e), despite that the responses were measured only in quiet conditions and in a normal healthy adult population. These results suggest that previously reported relationships may be present as a continuum in the population and even in optimal listening conditions. The MEG-FFR technique may allow us to more consistently observe behavioural and experience-related relationships with FFR-fO strength in less challenging listening conditions as compared with the EEG-FFR. The EEG-FFR is likely a composite from several subcortical [Chandrasekaran and Kraus, 2010] and cortical [Coffey et al., 2016b] sources. In recent work, we compared two common single-channel EEG montages (Cz-mastoids and Fz-C7) and found that while FFR-fO strength in each montage (measured simultaneously) was moderately correlated, a large proportion of variability was unaccounted for and the two methods differed in their sensitivity to a behavioural measure of interest [Coffey et al., 2016a]. This observation suggests that differences in individuals' head and brain geometry may sometimes obscure EEG-FFR vs behavioural relationships, possibly due to interference from source summation at a given point of measurement.

The spatial resolution of MEG source imaging may help to clarify the auditory processes that generate the ERP and ERF components, which have a long history in auditory neuroscience yet have predominantly been studied at the sensor level or using simpler models. We used distributed source modelling based on individual anatomy to localize the sources of each ERP/ERF wave (Figure 3d) with a view to confirming the localization of the wave of interest. The Pl wave originated bilaterally in the primary auditory areas, as expected [Key et al., 2005; Liegeois-Chauvel et al., 1994]. P2 appeared to be right-lateralized and comparatively more anterior along the superior temporal plane as compared to the Pl signal. This is generally consistent with previous work [Alain et al., 2013], but contrasts with an analysis of equivalent current dipoles that suggested a more posterior and medial source for the P2 [Shahin et al., 2003]. However, both the stimulation and analysis (e.g. use of the standard brain rather than individual anatomy) vary considerably between these studies, making this difference difficult to interpret.

The relatively more anterior location of the P2 compared to the FFR generators (Figure 4d) could be explained by a right-lateralized anterior flow of pitch-relevant information that supports SIN processing, although future work will be needed to clarify whether the relationship between periodic encoding and later waves is causal in nature, or if these different frequency bands represent neural activity in parallel processing streams in neighbouring neural populations. Future work could also aim to clarify how basic auditory information is separated and streamed to other cortical areas in order to accomplish different auditory tasks, using the spatial information in MEG data or a combinations of EEG and fMRI data.

The strength of signal generators in the right hemisphere was stronger for both the FFR-f0 and P2 waves (Figure 4d). We found a positive correlation between P2 amplitude (measured with EEG at Cz) and MEG FFR-f0 in the right hemisphere (but not left; Figure 4c,e). These results corroborate previous work suggesting that the right auditory cortex is relatively specialized for pitch and tonal processing [Albouy et al., 2013; Andoh et al., 2015; Cha et al., 2016; Herholz et al., 2015; Hyde et al., 2008; Mathys et al., 2010; Patel and Balaban, 2001; Patterson et al., 2002; Schneider et al., 2002; Zatorre et al., 1994]. It has been proposed that subtle differences in neural responses early in the cortical processing stream may lead to distinct functional roles for higher level processes out of a need for optimization, in particular a right-hemisphere bias for periodicity and left–hemisphere bias for fine temporal resolution [Zatorre et al., 2002]. The relatively stronger temporal representation of periodicity in the right hemisphere may be the underlying reason for which hemispheric asymmetry is observed in many cortical auditory phenomena [Coffey et al., 2016b]. The present results are congruent with these hypotheses.

Periodicity encoding is related to pitch information [Gockel et al., 2011], which is one of several cues that the brain can use to separate streams of auditory information [Moore and Gockel, 2002]. We previously showed that fine pitch discrimination skills are correlated with FFR-f0 strength in the right auditory cortex [Coffey et al., 2016b]. Here, we add that discrimination thresholds correlate with SIN accuracy, and that SIN accuracy is related to periodic encoding in the auditory cortex. Together, these results support a mechanistic explanation for SIN enhancement via better pitch processing leading to better stream segregation. This explanation can also account for musician advantages in SIN, as well as the inconsistency with which it is observed across studies, whose design may emphasize other SIN cues. In the subset of subjects who reported having had musical training, measures of the extent and timing of musical practice were related to behavioural measures of SIN accuracy and fine pitch discrimination, and were paralleled in physiology by relationships to FFR-f0 and P2 amplitude. Experience-dependent plasticity therefore likely tunes FFR-f0 strength and tracking ability [Bidelman, 2011; Carcagno and Plack, 2011; Musacchia et al., 2007; Song et al., 2008]. Stronger periodicity encoding might thereby account in part for a musician advantage. However, other top-down factors are also at play in SIN perception, including auditory working memory, long-term memory, and selective attention. Each of these may be influenced by experience, and other peripheral and central factors [Anderson et al., 2013b]. These latter factors would likely be related to top-down effects originating in extra-auditory cortical areas such as motor and frontal cortices [Du et al., 2014] whereas the feed-forward mechanisms we emphasize here likely represent neural modulations within ascending neural pathways including brainstem nuclei, thalamus, and cortex. One or both of these mechanisms may be enhanced by training.

We propose that whether a musician advantage in SIN perception is observed or not in a given study may depend on interactions between the current state of the auditory system and the specific cognitive demands of the SIN task used in the study. Specifically, performance can depend on 1) the cues offered to the listener in the SIN paradigm (e.g. spatial cues and degree of information masking; [Swaminathan et al., 2015]); 2) the degree to which an individual's experience has enhanced representations and mechanisms related to the available cues and caused them to be more strongly weighted; and 3) how well individuals can adapt to use alternative cues and mechanisms when one or more cues becomes less useful either through task differences like levels of noise (e.g. [Du et al., 2014]) or due to physiological deterioration [Anderson et al., 2013b].

## 5. Conclusion

In this study we present further evidence that the quality of basic feed-forward periodicity encoding is related to the clinically relevant problem of separating speech from noise signals, and musical training. Our results demonstrate that neural activity related to fine pitch processing in subcortical structures and areas of the right auditory cortex generates the frequency-following responses, and also slower, later cortical responses. Inter–individual differences in neural correlates of basic periodic sound representation observed within the normal–hearing population [Coffey et al., 2016a; Ruggles et al., 2011] may in part be responsible for the surprising variability in SIN perception observed in similar populations (e.g. [Swaminathan et al., 2015]). This work sketches in the spatial and temporal properties of a stream of pitch-relevant information from subcortical areas up to and beyond right primary auditory cortex. More work is needed to explore exactly how this information is routed and subsequently used by higher-level networks. We conclude that better sound encoding likely improves SIN perception through better stream segregation, which is one of several contributing processes to performance. Importantly, basic sound encoding is associated with training and experience, supporting efforts to develop training-based treatment strategies [Bidelman and Alain, 2015].

## Acknowledgements

We wish to thank Elizabeth Bock for assistance designing and testing the EEG-MEG recording set up, Marc Bouffard for help preparing subjects, Francois Tadel for his expert assistance with Brainstorm software, and Alexandre Lehmann for consultation regarding the interpretation of evoked auditory responses. The research was supported by operating grants to R.J.Z. from the Canadian Institutes of Health Research and from the Canada Fund for Innovation, by a Vanier Canada Graduate Scholarship to E.B.J.C. and by seed funding from the Centre for Research on Brain, Language and Music (CRBLM). S.B. was supported by the Killam Foundation, a Senior-Researcher grant from the Fonds de Recherche du Quebec-Sante, a Discovery Grant from the Natural Science and Engineering Research Council of Canada and the National Institutes of Health (2R01EB009048-05).

